# Peroxisome turnover and diurnal modulation of antioxidant activity in retinal pigment epithelia utilizes microtubule-associated protein 1 light chain 3B

**DOI:** 10.1101/687533

**Authors:** Lauren L. Daniele, Jennifer Caughey, Stefanie Volland, Rachel C. Sharp, Anuradha Dhingra, David S. Williams, Nancy J. Philp, Kathleen Boesze-Battaglia

**Author notes:** Corresponding Author Kathleen Boesze-Battaglia, PhD, Department of Biochemistry, SDM, University of Pennsylvania, Philadelphia, PA 19104.

## Abstract

The retinal pigment epithelium (RPE) supports the outer retina through essential roles in the retinoid the visual cycle, nutrient supply, ion exchange and waste removal. Each day the RPE removes the oldest ∼10% of photoreceptor outer segments through phagocytic uptake, which peaks in a synchronous burst following light onset. Impaired degradation of phagocytosed OS material by the RPE can lead to toxic accumulation of lipids, oxidative tissue damage, inflammation and cell death. OSs are rich in very long chain fatty acids which are preferentially catabolized in peroxisomes. Despite the importance of lipid degradation in RPE function, the regulation of peroxisome number and activity relative to diurnal OS ingestion is relatively unexplored. Using immunohistochemistry, immunoblotting and catalase activity assays, we investigated peroxisome abundance and activity at 6 am, 7 am (at lights on), 8 am, and 3 pm, in WT mice and mice lacking microtubule-associated protein 1 light chain 3B (LC3B), that have impaired degradation of phagosomes. We found that catalase activity, but not protein expression, is 50% higher in the morning compared with 3 pm, in RPE of WT but not *LC3B*^*-/-*^ mice. Surprisingly, we found that peroxisome abundance was stable during the day, however numbers are elevated overall in *LC3B*^*-/-*^ mice, implicating LC3B in autophagic organelle turnover in RPE. Our data suggest that RPE peroxisome function is regulated in coordination with phagocytosis, possibly through direct enzyme regulation, and may serve to prepare RPE peroxisomes for daily surges in ingested lipid-rich OS.

## Introduction

The retinal pigment epithelium (RPE) supports photoreceptor activity and outer-segment (OS) renewal through the retinoid visual cycle and ingestion of spent photoreceptor outer segments (36, 41, 42, 66, 73). On a daily basis, the RPE ingests 10% of the lipid and protein-rich OS, for recycling and degradation, while new OS material is added basally (69, 74). Incomplete phagosome degradation in the highly oxidative environment of the RPE predisposes undigested lipids to oxidative damage and the formation of lipid peroxidation adducts contributing over time to an immune response and tissue damage (3, 19, 27, 35, 70).

The ingested OS are a complex mix of saturated and highly unsaturated fatty acids, which make up 50% of OSs (by weight) and provide the RPE with substrate for mitochondrial and peroxisomal fatty acid oxidation (FAO) (1, 23, 61, 64, 68). Within the lipid pool, OS membranes are estimated to contain ∼ 30% very long chain fatty acids VLCFAs (23). VLCFAs (C≥20) are preferentially catabolized in peroxisomes via β-oxidation (64, 68). Therefore, efficient lipid catabolism in the RPE requires coordination of β-oxidation pathways within both mitochondria and peroxisomes (1, 23, 61). Once VLCFA are shortened, they are substrates for mitochondrial β-oxidation (1, 61). Some of the resultant acetyl-CoA is funneled into the mitochondrial ketogenesis pathway, producing beta hydroxybutyrate (βHB). βHB is subsequently secreted into the subretinal space and taken up by photoreceptors as a metabolic substrate (1). Given their intimate metabolic coordination it is not surprising that numerous micro-peroxisomes can be found in RPE cells where they are contiguous with ER tubules and proximal to mitochondria (45, 47, 62).

Peroxisomes are ubiquitous organelles highly enriched in liver, intestine, adipose tissue and brain. In addition to their functions in α and β oxidation-mediated catabolism of fatty acids (48) peroxisomes play essential roles in ether lipid synthesis, docosahexaenoic acid (DHA) synthesis, and bile acid synthesis. Peroxisomes are also sites for both the generation and reduction of reactive oxygen and nitrogen species (ROS and RNS) which serve as mediators for cellular signaling (56, 72). The most abundant of these is hydrogen peroxide, produced as a byproduct of the initial desaturation of Acyl-CoA esters by Acyl CoA Oxidases (ACOXs) in the committed step of peroxisome fatty acid oxidation (64, 68). Hydrogen peroxide generated in the ACOX reaction is rapidly degraded by the antioxidant enzyme, catalase. Peroxisome antioxidant activity declines with cellular age (44). Dysregulation of peroxisome antioxidant capacity with age may have broad cellular impact as inhibition of catalase activity in fibroblasts resulted in increased mitochondrial ROS levels and disrupted activity (44, 67, 71). Catalase activity is also lower in peroxisomes of human primary RPE from aged donors, and those with age-related macular degeneration (AMD) (46), and paradoxically, this is accompanied by an overabundance of peroxisomes (6, 7, 21), suggesting a role for impaired peroxisome turnover in declining peroxisome function.

Although countless studies have focused on understanding peroxisome proliferator-activated receptor (PPAR) regulation, and correlation with disease (50, 60, 63, 76), to our knowledge little information is available regarding the activity and numbers of peroxisomes in the RPE over a 12hr light/dark cycle. In this study, we investigate peroxisome function and dynamics in the context of phagocytic degradative processes in RPE cells. We find that catalase activity is increased by ∼50% during the time of peak phagocytosis, and this diurnal regulation of antioxidant activity is absent in mice lacking LC3B. We also find that while LC3B-dependent processes contribute to peroxisome turnover, peroxisome number does not depend on time of day. Our data suggest peroxisome function in the RPE is regulated dynamically in coordination with phagocytosis, likely through direct regulation of enzyme activity. Catalase activity can be regulated through diverse post-translational modifications (9, 10, 16, 26, 39, 59) which are often sensitive to the redox state of the cell. Elevated catalase activity during the early morning suggests the possibility of a homeostatic mechanism to ready RPE peroxisomes for the daily synchronous surge in ingested lipid-rich outer segments.

## Results

### Mouse RPE contain abundant peroxisomes

Fatty acids comprised of 20 carbons and longer are catabolized in peroxisomes (64, 68), where the rate limiting step is desaturation of a fatty acyl-CoA by the enzyme ACOX1, producing the byproduct hydrogen peroxide (H_2_O_2_) (Fig.1A). Catalase, the most abundant peroxisomal antioxidant, reduces H_2_O_2_ produced in peroxisomes during this reaction, preventing its toxic accumulation. Using a publicly available microarray data set (GSE10246; Fig 1B) we analyzed gene expression profiles of peroxisome lipid-associated proteins in mouse tissues. RPE express fatty acid transporters and peroxisome β-oxidation enzymes. Three members of the ABC lipid-transporter superfamily reside in peroxisomes, these include, *Abcd1* (ALDP), *Abcd2* (ALDR) and *Abcd3* (PMP70). Of these, *Abcd1* is expressed in RPE (mouse) in relative abundance with levels of *Abcd3* in RPE similar to that in liver (Fig. 1B). Transcripts for *Acox1*, and catalase as well as 3-ketoacyl-CoA thiolase (*Acaa1/*thiolase) are highly expressed in liver but are also enriched in the RPE relative to retina, cornea and neural tissues (Fig 1A, B). Two thiolase genes are expressed in mouse, *Acaa1a*, and *Acaa1b*, with the expression of thiolase b restricted to liver, with lower expression in kidney, intestine and adipose tissue (13). The complete β-oxidation of polyunsaturated fatty acids (FA) within peroxisomes requires additional enzymes, due to the presence of *cis-* and *trans-* double bonds (68), including peroxisome 2,4, Di-enoyl CoA reductase (*Decr2*). We find that transcripts for these enzymes are also enriched in RPE relative to retina and brain <u>(Supplemental Fig. S1)</u>.

**Figure 1.**
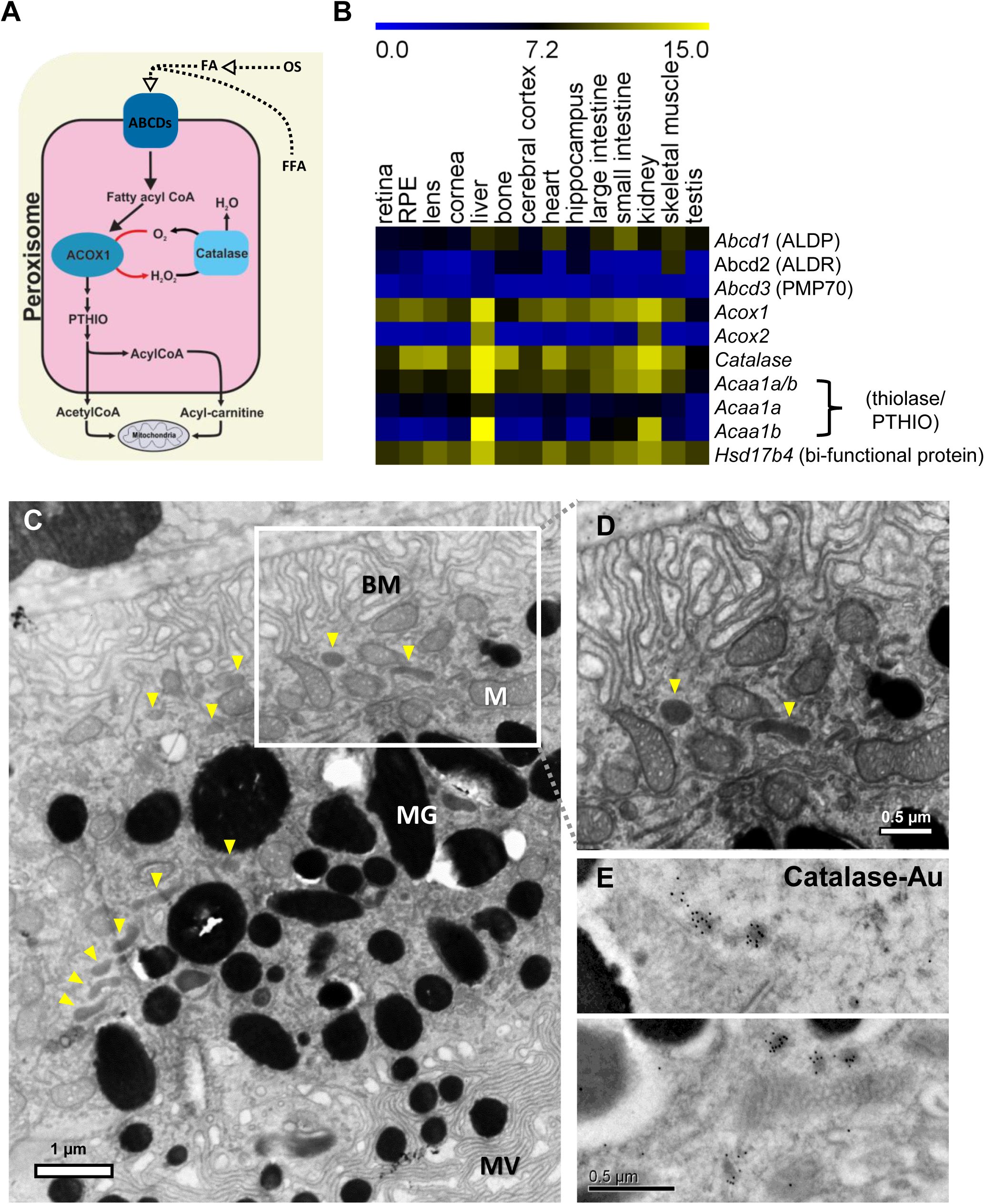
Enrichment of β-oxidation genes in peroxisomes of mouse RPE and their identification with TEM. A. Schematic depicting key enzymes involved in fatty acid β-oxidation within peroxisomes. The initial committed step, desaturation of acyl - CoA esters by ACOX1, produces H_2_O_2_ as a byproduct, which is then neutralized by catalase. FFA – free fatty acid, OS – photoreceptor outer segments. B. Microarray expression analysis from GSE10246 showing relative enrichment of peroxisomal β oxidation enzymes, including catalase, in mouse RPE. C. Transmission EM image of WT RPE labeled with DAB. BM - RPE basement membrane, M-mitochondria, MG – melanin granule, MV - microvilli. D. Boxed field (in panel C) at higher magnification. Yellow arrowheads in C and D indicate peroxisomes. E. Example images of Immuno-electron microscopy in WT mouse RPE showing gold particle labeling of catalase.

We next examined peroxisome localization and abundance in the RPE, using 3,3’-diaminobenzidine (DAB) reactivity by transmission EM complemented with immuno-electron microscopy analysis. Catalase reacts with DAB in the presence of excess H_2_O_2_ to form a dense precipitate localized to catalase-positive peroxisomes (15). DAB was localized to small oval shaped organelles with ∼100 nm diameter in ultrathin sections of RPE (Fig. 1C, D). When Immuno-electron microscopy (EM) analysis was used to detect catalase, gold particles labeled small rounded structures adjacent to mitochondria (Fig 1E), morphologically consistent with our DAB labeling. In RPE whole mounts labeled with PMP70 and Pex14, we observe numerous peroxisomes throughout the cytoplasm, some with a tubular morphology <u>(Supplemental Fig. S2)</u> as was recently observed for peroxisomes of mouse fibroblasts using stimulated emission depletion (STED) microscopy (65). Catalase immunolabeling appeared most intense in the RPE relative to other retina layers in cryosections of mouse eye (Fig 2 A). In RPE, catalase-positive puncta were enriched in the peroxisome-specific membrane protein, PMP70, a fatty acid ABC transporter used as a marker of peroxisome number (22, 51) (Fig 2B,C). Linear intensity profiles show that peaks in catalase signal are spatially correlated with that of PMP70 (Fig 2C) and co-distribution analysis of catalase and PMP70 have a Pearson’s correlation coefficient of 0.89 (± 0.006 n=38). We also examined catalase expression in RPE of *microtubule-associated protein 1 light chain 3B*^*-/-*^ *(LC3B*^*-/-*^) mouse, a model of defective phagosome maturation and lipid dysregulation (18, 19). *LC3B*^*-/-*^, like WT also show a relative enrichment of catalase in RPE relative to retina (Fig 2 D), with linear intensity profiles of PMP70 and catalase well correlated (Fig 2 E,F). Our data shows that relative to other cells within the eye, peroxisomes are abundant in RPE, with catalase co-distributing with PMP70-positive peroxisomes in both *LC3B*^*-/-*^ and WT mice.

**Figure 2.**
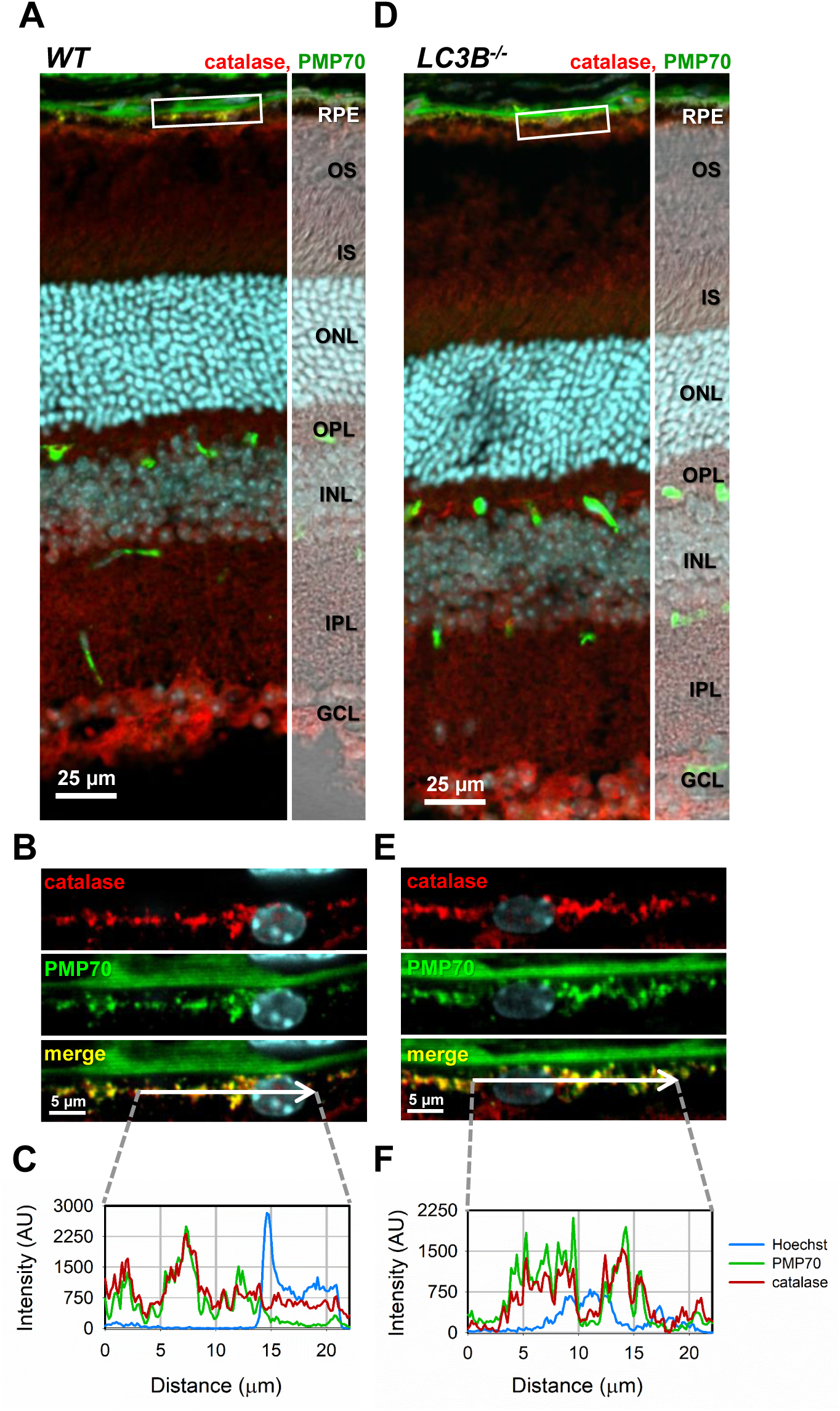
Mouse RPE peroxisomes are enriched in catalase. A and D. Single plane confocal micrographs of cryosections of WT and *LC3B*^*-/-*^ mouse eyes harvested and fixed at 10 am and immunolabeled with antibodies to PMP70 (green) and catalase (red). GCL - ganglion cell layer, IPL-inner plexiform layer, INL – inner nuclear layer, OPL-outer plexiform layer, IS – photoreceptor inner segments, OS-photoreceptor outer segments, RPE - retinal pigment epithelium. B, E. Higher magnification views of the boxed regions in A and D respectively. C, F. Intensity profiles for PMP70 (green), Catalase (red) and Hoechst nuclear dye (blue) across the lines depicted in B, and E, respectively. The degree of overlap of catalase and PMP70 was estimated with a Pearson’s correlation coefficient of 0.89 (± 0.006 n=58) for WT and 0.89 (± 0.005 n=55) for *LC3B*^*-/-*^ measured from circular ROIs (5 μm diam).

### Peroxisome numbers do not depend on time of day

Peroxisomes respond to changes in the cellular environment by adapting their number, morphology and metabolic functions. In RPE, the level of VLCFA substrate for peroxisome oxidation is regulated in a synchronous circadian manner through phagocytosis of OSs (28, 41, 42). Phagocytic ingestion of rod photoreceptor OS peaks following light onset (42). Because peroxisome number can be dynamically regulated by cellular conditions, we wanted to determine if OS disc phagocytosis influences peroxisome abundance and function in the RPE. We analyzed peroxisome abundance in WT mouse RPE at various times of day and compared this to peroxisome abundance in the RPE of the *LC3B*^*-/-*^ mouse, having defective phagosome maturation (18, 19). RPE/choroid was isolated at: 6 am - one hour before lights on, 7 am – at lights on, 8 am - an hour after lights on, and 3 pm (8h after light onset). We compared the levels of two membrane proteins widely used as markers for peroxisomes; Pex14 is an essential transmembrane component of the peroxisome translocation machinery that imports cytosol-translated enzymes into the peroxisome lumen (77) and PMP70, one of 3 ATP binding cassette family members that transport FAs into peroxisomes (22, 51). Pex14 (Fig 3 A, C) and PMP70 levels (Fig 3 B, D) remained relatively stable throughout the day in RPE/choroid isolated from WT mice. When compared to WT mice, the levels of Pex14 in the *LC3B*^*-/-*^ mouse RPE were elevated by 20% of WT at 7 am and 12% at 8 am, whereas PMP70 expression was elevated above WT by an average of 32% at all time points (Fig 3 E, F). Peroxisome abundance was next assessed in *LC3B*^*-/-*^ and WT mouse RPE by confocal imaging (Fig.4 A, B). Pex14 immunoreactivity increased by at least 50% in *LC3B*^*-/-*^ vs WT in 2 data sets (Fig 4C). PMP70 immunoreactivity was increased by 50% in one data set with another data set showing a trend towards higher PMP70 immunoreactivity that was not statistically significant. A third data set showed no difference between *LC3B*^*-/-*^ and WT immunoreactivities. Collectively, these studies suggest that peroxisome degradation is reduced in RPE of *LC3B*^*-/-*^ mice.

**Figure 3.**
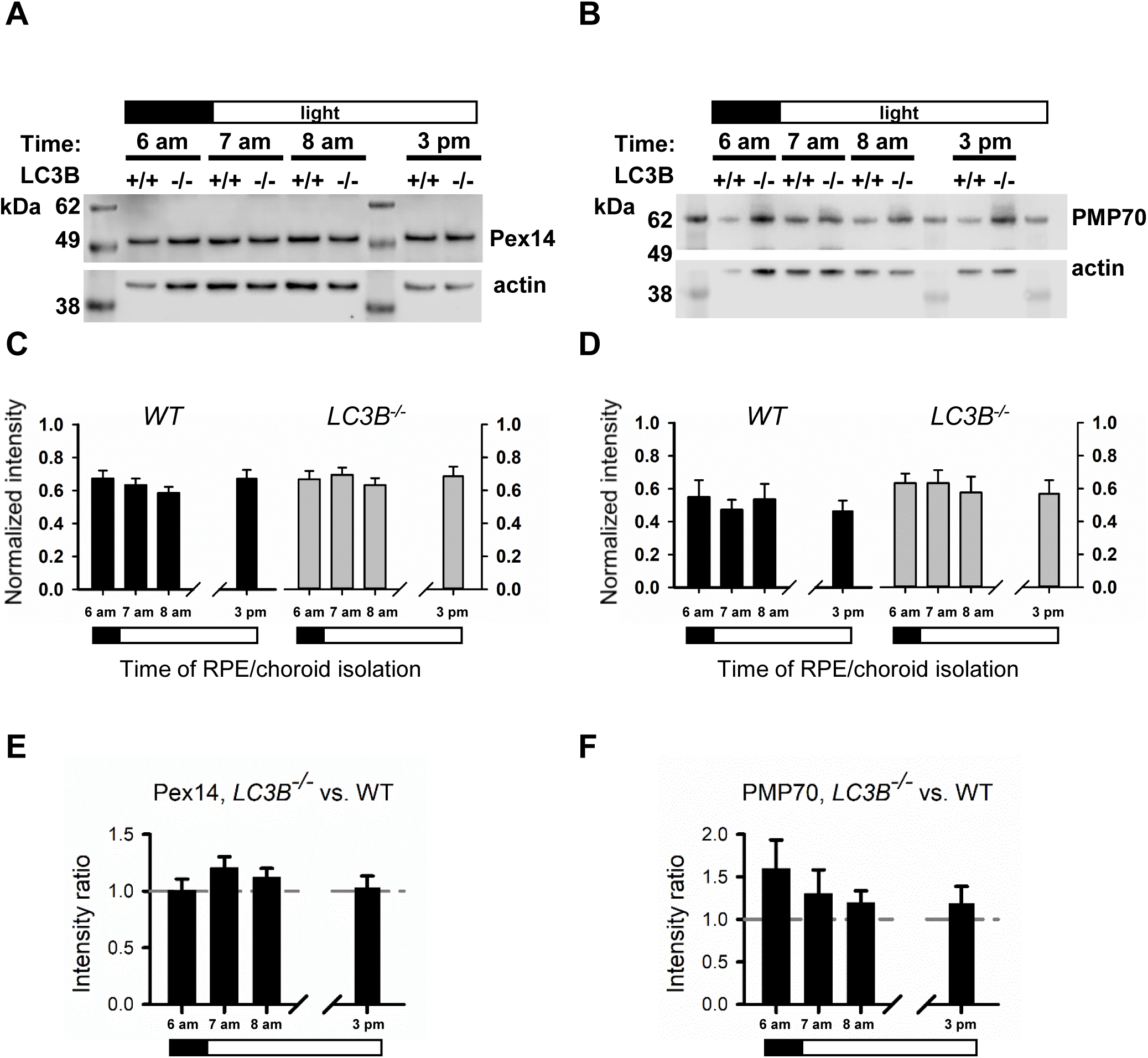
Expression of Pex14 and Pmp70 is elevated in *LC3B*^*-/-*^ during the day but does not vary diurnally. A, B. Western blots of RPE lysates made at indicated times of day. Immunoblotting was performed with antibodies to peroxisomal membrane proteins Pex14 (A) and PMP70 (B) and loading control actin. C, D. Quantification of intensities of Pex14 (C) or PMP70 (D) relative to loading control band intensities as a function of time of day for WT (left plots) and *LC3B*^*-/-*^ lysates (right plots each panel). Values were not significantly different in C and D. Ratio of immunoreactivity of Pex14 (E) or PMP70 (F) in *LC3B*^*-/-*^ vs. WT estimated from the same data in C and D, respectively. Values were not significantly different across each time point in E and F.

**Figure 4.**
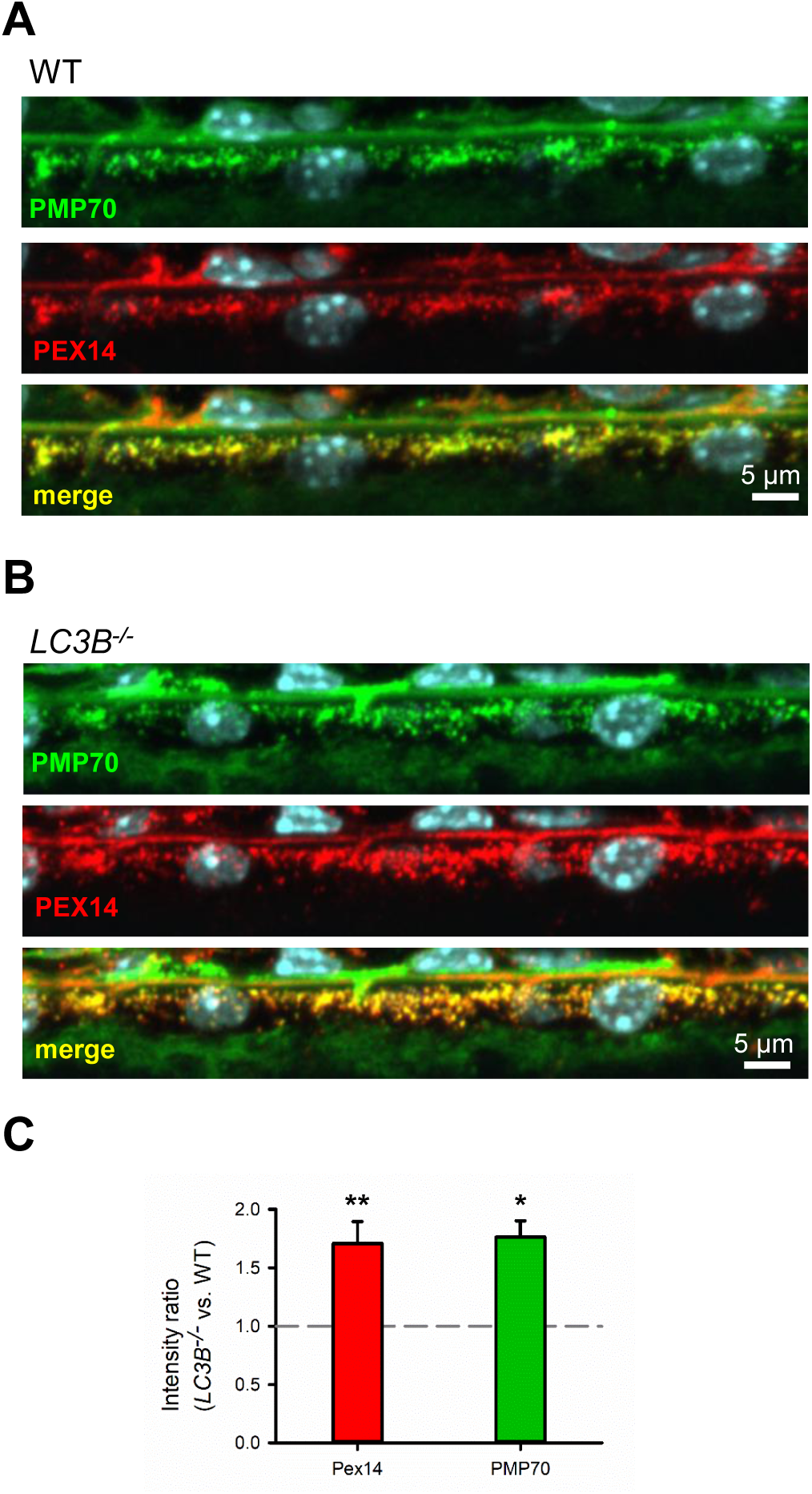
Peroxisomes are more numerous in *LC3B*^*-/-*^ mice. A, B. Cryosections of eyes from WT and *LC3B*^*-/-*^ mice harvested at 10 am showing the RPE layer, and immunolabeled with antibodies to peroxisome specific markers Pex14 (red) and PMP70 (green) Hoechst nuclear dye (cyan). C. Example ratios of fluorescence intensities for Pex14 or PMP70 in *LC3B*^*-/-*^ vs. WT images, asterisks denote statistical significance between WT and *LC3B*^*-/-*^ intensities: ** p < 0.01, * p < 0.02.

### LC3B contributes to early morning peak in catalase activity in RPE

Although peroxisome numbers are relatively constant over the time period tested, because RPE ingests a lipid rich OS at light onset, we sought to determine if peroxisome enzymatic activity depends on substrate availability. We compared ACOX1 activity as coupled catalase function in WT and *LC3B*^*-/-*^ mice. Therefore, we compared peroxisomal catalase activity at 6 am, 7 am (at lights on) and 8 am – during which the OS-derived LCFA and VLCFA β-oxidation substrates are maximal- and at 3 pm, 8 hours later. At the 8hr time point virtually no phagosomes were observed in WT mice while over 50% of the phagosomes remain undegraded in the *LC3B*^*-/-*^ mouse RPE (18, 19). Catalase activity was elevated ∼1.5 fold during the early morning compared with the afternoon in WT mice (Fig 5 A), but not in *LC3B*^*-/-*^ mice, suggesting that an LC3B dependent process influenced peroxisome catalase activity. To determine if elevated catalase activity was due to increased enzyme levels, we investigated catalase protein amounts from the same mice at 6 am, 7 am (at lights on), 8 and at 3 pm, by immunoblot analysis. Catalase protein levels remained relatively constant over the time periods studied (Fig. 5B, C). In contrast to our catalase activity data, we observed an apparent increase in relative catalase protein levels between 8 am and 3 pm in WT, however catalase protein levels at 6 am and 7 am were not significantly different from those at 3 pm. We quantified catalase immunoreactivity from our immune-EM analysis of WT mouse eyes removed at 1 hour and 6 hours after light onset. At 1 hour after light onset, we found 467 gold particles / μm^2^ (± 30 sem, n=41), which was comparable to our finding of 432 particles/ μm^2^ (± 20 sem, n=39) at 6 hours after light onset. Taken together, our data suggest that changes in catalase protein levels do not account for the elevated activity during the morning. We also tested whether diurnal modulation of metabolic activity was unique to peroxisomes, and so we investigated whether mitochondrial activity was also regulated diurnally by assessing activity of citrate synthase, the first enzyme of the tricarboxylic acid (TCA) cycle (Fig 5D), and found no difference in citrate synthase activity as a function of time of day. We directly tested whether catalase is activated by increased VLCFA and LCFA β-oxidation substrate after uptake of OSs or by elevated H_2_O_2_, by measuring catalase activity in RPE/choroid explants treated with OSs or H_2_O_2_. Catalase activity was increased by ∼50% above Ringers control (p<0.05) in RPE explants incubated with 0.5 mM H_2_O_2_ but not OSs (Fig 5E). We found no statistically significant differences in catalase activity in RPE explants from *LC3B*^*-/-*^ for either treatment.

**Figure 5.**
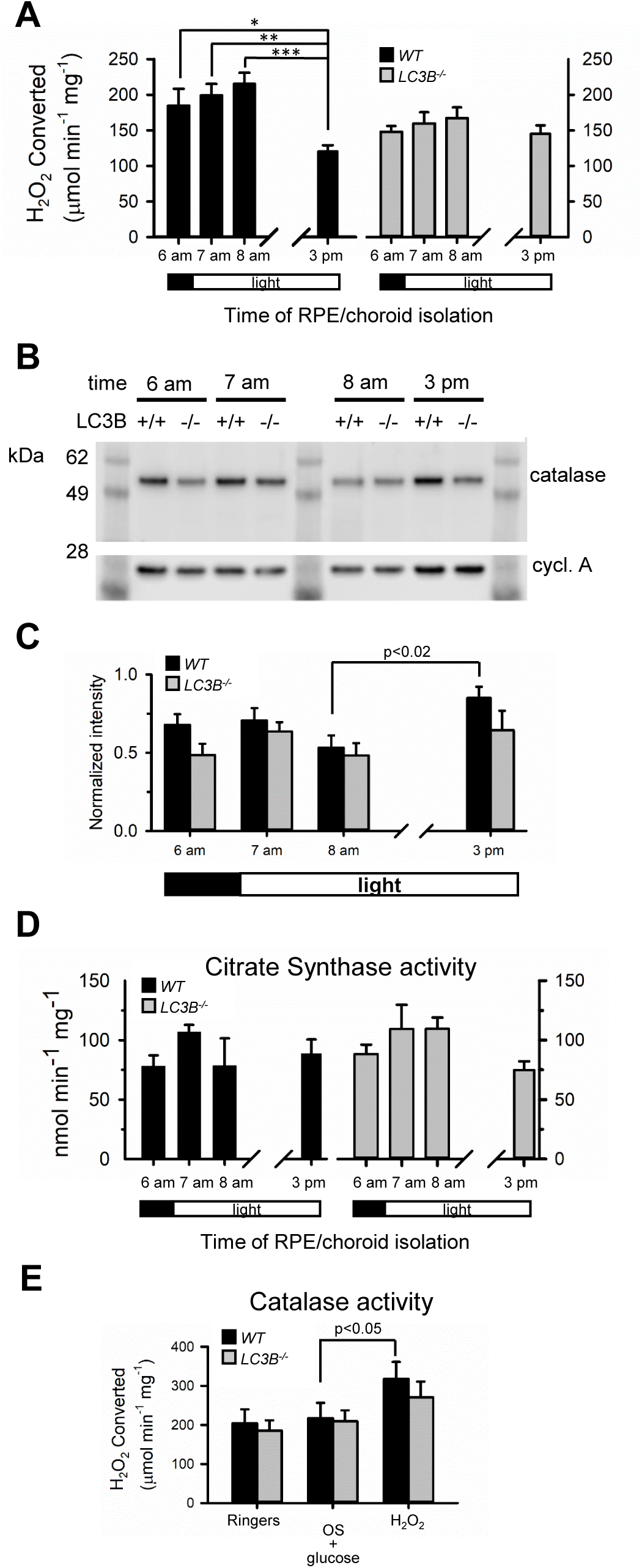
Elevated catalase activity during early morning in RPE of *WT* but not *LC3B*^*-/-*^. A. Catalase activity measured from RPE/choroid lysates made at different times relative to light onset (7 am). Activity is expressed relative to mg protein. 8 data sets from 3 mice of each genotype were averaged. *p<0.02, **p<0.0006, **p<0.0001. Values were not significantly different across time points for *LC3B*^*-/-*^ mice (one-way ANOVA). Differences in catalase activities between WT and *LC3B*^*-/-*^ were statistically significant (p<0.009 two-way ANOVA), but at 3 pm, WT and *LC3B*^*-/-*^ catalase activity differences did not reach statistical significance (p<0.1). B. RPE choroid lysates from WT and *LC3B*^*-/-*^ mice made at the indicated times of day were immunoblotted with antibodies to catalase, and cyclophilin A (as a loading control). C. Relative intensities of catalase vs. cyclophilin A in immunolabeled bands from Western blots of RPE/choroid lysates made at different times of day. D. Citrate synthase activity in WT and *LC3B*^*-/-*^ lysates of posterior eyecup after removal of neural retina (RPE/choroid). Values were not significantly different. E. Catalase activity in lysates of RPE/choroid explants incubated with Ringers, or with added bovine outer segments and 5 mM glucose, or 0.5 mM H_2_O_2_. Catalase activity was increased by 55% in WT after H_2_O_2_ treatment compared with Ringers control (*p<0.05).

## Discussion

### RPE peroxisomes are enriched in β-oxidation genes

Peroxisomal β-oxidation preferentially breaks down very long chain fatty acids abundant in POS membranes. The chain-shortened FAs can then be utilized by mitochondria for further rounds of β-oxidation. In this study we show that genes encoding key enzymes of fatty acid β-oxidation within peroxisomes, such as Acox1, and catalase are highly expressed in mouse RPE relative to other eye tissues. Peroxisome β-oxidation genes are most highly expressed in liver, consistent with the liver’s principal role in fatty acid metabolism and regulation of systemic fatty acid availability (48, 53). Peroxisomes in the liver, unlike adipocytes, brain and intestinal mucosa, are large and contain a matrix enriched in luminal enzymes, including catalase (48). β-oxidation pathways of the liver play multifaceted roles in fatty acid metabolism, including the breakdown of dicarboxylic acids and dietary pristanoyl-CoA. Single rounds of β-oxidation in liver are also critical for synthesis of bile acids from cholesterol and DHA from essential fatty acids (5, 68). Previous studies have shown that RPE like the liver, is ketogenic, with the ability to catabolize fatty acids (FAs) derived from phagocytosed photoreceptor outer segments (POS) through mitochondrial β-oxidation pathways that produce acetyl-CoA and the secreted ketone metabolite β-hydroxybutyrate (1, 61). Peroxisomes of the RPE are found adjacent to mitochondria and their proximity supports their coordination of metabolic pathways. In addition to metabolic coordination, mitochondria and peroxisomes overlap in redox sensing and ROS/RNS signaling and regulation. Mitochondria and peroxisomes both produce significant amounts of ROS through their different metabolic pathways. Each round of β-oxidation in peroxisomes produces H_2_O_2_. We show that the antioxidant catalase, the primary enzyme responsible for the removal of H_2_O_2_ is enriched in peroxisomes within RPE. Peroxisomal antioxidant activity, plays a key role in redox homeostasis, and disrupted catalase activity has serious consequences for mitochondrial health (44, 67, 71).

### Catalase activity may be regulated post-translationally

We show that catalase, the heme-containing antioxidant enzyme enriched in RPE peroxisomes, has highest activity during the early morning when OS derived VLCFA’s are expected to reach their maximum in RPE cells and β-oxidation reaches its peak (61). The elevated catalase activity is not due to increased expression of catalase protein, suggesting that diurnal modulation of catalase activity occurs at the post-translational level. Catalase activity is regulated by a diverse array of post-translational modifications depending on cell context, including phosphorylation, S-nitrosylation, oxidation and ubiquitination (9, 10, 16, 26, 39, 59), as well as regulation of its multimerization (59). Catalase is thought to exist mainly in a tetrameric, enzymatically active form, but higher-order oligomer and dimer forms have been encountered inside cells (2, 58). The enzymatic activities of catalase in H_2_O_2_ removal as well as its peroxidase activity, requires its heme group and heme binding is required for tetramerization and maturation (11). During small elevations in cellular H_2_O_2_ in cultured mouse fibroblasts, the redox sensitive kinases, c-Abl and Arg, increase catalase activity by phosphorylation of key tyrosines and directly binding, protecting catalase from de-phosphorylation (9, 10). (59) Catalase is an important antioxidant whose activity declines with age and in AMD-afflicted RPE (46) and the regulation of its activity in RPE is largely unexplored.

### Diurnal modulation of Catalase activity depends on LC3B

In contrast to WT RPE, RPE of *LC3B*^*-/-*^ mice lack a boost in catalase activity during the morning. RPE of *LC3B*^*-/-*^ mice have impaired degradation of phagocytosed OS – derived material, depressed fatty acid oxidation, eventual accumulation of lipid deposits, and low levels of esterified and free DHA, due in part to the role of LC3B in LC3-associated phagocytosis (19). Lower catalase activity in RPE of *LC3B*^*-/-*^ mice during the morning is consistent with impaired β-oxidation of POS-derived FAs in *LC3B*^*-/-*^ (19). Elevated catalase activity in RPE may reflect higher peroxisomal β-oxidation activity during the morning when VLCFA availability may be maximal. The synchronous burst in POS uptake by phagocytosis in mammalian RPE occurring ∼1 hour after light onset is necessary for the maintenance of photoreceptor health (52). We hypothesized that catalase activity is modulated in step with this burst, through increased H_2_O_2_ produced when availability of OS – derived LCFA and VLCFA substrate reaches its peak. If this were true, RPE explants made from WT mice fed OS may respond with a boost in catalase activity, and this may be impaired in RPE explants from *LC3B*^*-/-*^ mice. However, increasing H_2_O_2_ independently of substrate availability may override the impairment and act more directly on a modifier of catalase activity (e.g. an H_2_O_2_-sensitive kinase) resulting in increased catalase activity in both genotypes. Our results show that H_2_O_2_ resulted in ∼50% increase in catalase activity for both WT and *LC3B*^*-/-*^ RPE explants, making H_2_O_2_ - sensitive kinase mediated phosphorylation a plausible mechanism in RPE cells. However, elevated catalase activity was not observed after treatment of RPE explants of either WT or *LC3B*^*-/-*^ mouse with OS, leaving open the possibility of catalase activity modulation through another pathway which is impaired in the absence of LC3B. Future studies are needed to decipher the role of LC3B in modulating catalase activity.

### RPE peroxisome number does not vary with time of day

Peroxisome number is dynamically regulated to adapt to the changing cellular milieu. The number of peroxisomes at any given time emerges from the dynamic balance of biogenesis, replication, and degradation (14, 30). In this study we explored the possibility that peroxisome number is dynamically regulated after the synchronous peak in outer segment phagocytosis, when ingested very-long chain fatty acids derived from photoreceptor outer segments, is maximal. However, our immunoblot evidence suggests RPE peroxisome mass is relatively stable over time. This was a surprise as previous studies in mouse show 2 brief peaks in RPE peroxisome number 3 hours before and 3 hours after light onset (47), suggesting rapid expansion followed by rapid removal of peroxisomes in RPE. The chronology of the observed peak in peroxisome numbers by EM roughly corresponds to our observations of peak catalase activity in the morning spanning an hour before and after light onset. However, we saw no evidence for rapid removal of peroxisome material. It is likely that peroxisome removal and replacement is a relatively slow process, as pulse-chase studies of fluorescently-tagged or isotope-labeled peroxisomal proteins indicated a half-life of 1-3 days for liver peroxisomes (31, 57). The observations of a steep decline in peroxisome numbers 1-2 hours after light onset track a peak in numbers of phagosomes in this study, suggesting the possibility of false negatives or misclassification of structures, when phagosomes accumulate and traffic more basally.

### Peroxisome degradation depends on LC3B; implications for RPE homeostasis

Turnover is important in maintaining healthy functional organelles. Degradation of dysfunctional and damaged peroxisomes and other organelles is principally achieved through regulated macroautophagy (or pexophagy) a lysosomal degradative process whereby a double membrane autophagosome engulfs the entire organelle prior to fusion with lysosomes (4, 34). The signals which select old dysfunctional peroxisomes for degradation are beginning to be elucidated and selective ubiquitination plays an important role (37). Prolonged residency of monubuqitinated Pex5 at the peroxisomal membrane is a signal for binding of pexophagy adapters P62 and NBR1 (17, 55). P62 contains an LC3 interaction motif and ubiquitin binding domains allowing its co-recruitment of peroxisome cargo and autophagosome machinery. In a distinct pexophagy pathway, under conditions of elevated ROS, ataxia-telangiectasia mutated kinase phosphorylates Pex5, allowing monoubiquitination at a lysine residue, leading to recruitment of P62 and autophagosomal membranes (75). Our results show that peroxisomes are more numerous in RPE of *LC3B*^*-/-*^ mice, which likely reflects sluggish degradation and turnover of peroxisomes, given the well-established role of LC3B in autophagosome formation and autophagosome-lysosome fusion (54). LC3B is one of 6 different members of the mammalian Atg8 family (LC3/GABARAP proteins) involved in autophagy (43). Of the LC3 subfamily, only LC3A and LC3B are expressed in mouse RPE (18). Our data suggests that the LC3B isoform is necessary for efficient turnover of peroxisomes within the RPE. This is supported by evidence of elevated peroxisome membrane proteins (Pex14 and PMP70) in RPE of *LC3B*^*-/-*^ mice from western blot and IHC. Our data is consistent with recent finding that LC3B was necessary for autophagic degradation of p62, normally degraded along with its cargo, in HEK293T cells (49).

In this study we found that RPE of mice lacking LC3B expression have elevated peroxisome numbers but lack the ability to increase activity of peroxisome antioxidant catalase during the early morning. Elevated peroxisome numbers with lower antioxidant function is reminiscent of aged and diseased RPE (6, 21). Reduced autophagic flux, accompanied by accumulation of autophagosomes and lysosomes, in RPE derived from AMD patients was proposed to contribute to poor mitophagy and mitochondrial dysfunction (29). The role for autophagic degradation in mitochondrial quality control in AMD has been the subject of more intense focus (32). Mitochondria and peroxisomes intimately cooperate in metabolic pathways and regulation of ROS. Impairments in autophagy in ageing and diseased RPE may have more far reaching consequences when the health and function of peroxisomes is also considered. Peroxisome turnover defects may impact the resilience of RPE to oxidative stress in ageing and contribute to disease pathogenesis.

## Materials and Methods

### Animals

C57BL6/J (WT) mice and the *LC3B*^*-/-*^ mouse line (strain name: B6;129P2-Map1Lc3b^tm1Mrab^/J; stock # 009336, (8) were purchased from Jackson Laboratory (Bar Harbor, ME). The *LC3B*^*-/-*^ mice were backcrossed for at least 5 generations onto a C57BL6/J background. The *LC3B*^*-/-*^ and the WT mice were confirmed to be free of the rd8 mutation by genomic DNA PCR using primers as described in (12). Maintenance of mouse colonies and all experiments involving animals were as described previously (24, 25). Mice were housed under standard cyclic light conditions: 12-h light/12-h dark and fed ad libitum, with both female and male mice used in these studies. All procedures involving animals were approved by the Institutional Animal Care and Use Committees of the University of Pennsylvania and the University of California, Los Angeles, and were performed in accordance with the Association for Research in Vision and Ophthalmology guidelines for use of animals in research.

### Microarray analysis for heat map

The heat map was generated using the publicly available high throughput *Mus musculus* gene expression microarray data set GSE10246 (40) available through NCBI GEO (GNF Mouse GeneAtlas V3). The probe IDs for each gene of interest were obtained from an up to date microarray annotation file (Mouse430_2na36annot) obtained from the Affymetrix website and used to search GSE10246. The heat map was made in MultiExperiment Viewer version 4.9.0 using Log2 scale for values averaged from 2 replicates for each tissue.

### Immunohistochemistry (IHC)

Eyes were removed from euthanized mice and immediately placed in 4% paraformaldehyde (PFA). For cryosections, the anterior segment and lens were removed during fixation, leaving an intact posterior eye (eyecup). Eyecups were fixed overnight at 4°C. Following cyroprotection in 30% sucrose, 1x PBS at 4°C overnight, eyecups were embedded in OCT compound (Sakura Finetek, Torrance, CA), frozen, and sectioned with a cryostat (Microm HM 550) at 10-20 μm thickness. Cryosections were rehydrated and washed in 1x PBS followed by incubation with antibodies diluted in blocking buffer (5% BSA, 0.1% triton X-100, 1X PBS). Sections were washed 3x in 1x PBS and bound primary antibodies were detected by incubation with Alexa-fluor conjugated secondary antibodies diluted in blocking buffer (Thermo Fisher/Invitrogen, Eugene, OR) for 1 hr at 37°C. For flat mounts of RPE/choroid/sclera, melanin and other pigment was bleached after fixation, using the method of Kim and Assawachananont (38). Briefly, each eyecup was incubated at 55° C in 10% peroxide/1x PBS for 2.5 hr and immunolabeled as above after 3-4 washes in PBS. Immunolabeled tissue was imaged with a Nikon A1 confocal microscope (Nikon Instruments, Melville, NY). We used the following primary antibodies for IHC (with company and dilution): rabbit anti-catalase (Abcam ab1877, 1/500), rabbit anti-Pex14 (ProteinTech 10594-1-AP, 1/500), mouse anti-PMP70 (Sigma SAB4200181, 1/300).

### Electron microscopy with DAB labeling of peroxisomes

Peroxisomes were labeled with alkaline 3, 3′-diaminobenzidine (DAB) following established methods (20). Briefly, mouse eyes were enucleated, and anterior segment removed as above, and immersed in fixative (2.5% glutaraldehyde, 2% paraformaldehyde, in 0.1M cacodylate buffer) overnight at 4°C. After fixation, small pieces of eyecup were washed 2x 30 min in 0.1M Cacodylate buffer, then washed 20 min in Tris HCl, pH 8.5, then in 0.2% DAB for 0.5 hr in darkness in a water bath at 37°C. Control tissue was incubated at 37°C with Tris HCl buffer alone. DAB solution contained 0.2 DAB (from a 4% stock; EMS 13080), 0.1 M Tris HCl, 0.15% H_2_O_2_. Following DAB incubation, tissue was washed 15 min 0.1M Tris HCl in darkness, then 15 min 0.1M Cacodylate in darkness. The labeled tissue was embedded in Epon. All post-fixation, embedding and ultrathin sectioning was performed by the University of Pennsylvania EM Resource Laboratory. Ultrathin sections were imaged with a Jeol-1010 transmission electron microscope.

### Immuno EM

Mouse eyes were enucleated and fixed at 4°C in 4% formaldehyde and 0.2% glutaraldehyde in 0.1 M sodium cacodylate buffer. The anterior segment was removed, and eyecups were dissected along the dorsal ventral axis. After washing in 0.1 M sodium cacodylate buffer, samples were dehydrated with an ethanol series, and embedded in LR-white resin. Ultrathin sections were collected on formvar coated nickel mesh grids, quenched with 0.1% glycine in 0.1M phosphate buffer for 15 min, blocked in 2% BSA in 0.1M phosphate buffer for 30 min, and incubated at 4°C overnight, with catalase ab15834 (Abcam, Cambridge UK) primary antibody, diluted by 1:500. Sections were then washed in 0.1M phosphate buffer, incubated with anti-rabbit IgG conjugated to 18-nm gold particles (Jackson Immuno Research Labs, West Grove, PA), washed again, and stained with 5% uranyl acetate in ethanol for 5 min. All samples were imaged with a JEM 1200-EX (JEOL, Peabody, MA) at 80kV at magnifications of 30,000 to 60,000x. Labeling density of the peroxisomes was determined by counting the number of gold particles per μm2 of sectioned peroxisome. Measurements were carried out using Fiji (ImageJ Version2.0.0; image processing software package; available at https://fiji.sc/). Data were collected from animals fixed at 1 and 6 hours after light onset.

### RPE/Choroid explants, OS and peroxide treatment

RPE explants were incubated in bicarbonate buffered Ringers with L-carnitine, or Ringers and either bovine OS particles (100,000 per chamber) or peroxide (0.5 mM) using published methods (61). Bovine outer segments were obtained from InVision BioResources (Seattle, WA). Stocks of 2 x 10^7^ particles/ml were made and stored in 15% sucrose. OS or peroxide were diluted to their final concentrations in Ringer’s and explants were incubated for 1-2 hrs at 37C with 5% CO_2_/O_2_.

### Western blotting

RPE explant lysates were prepared as described (Reyes-Reveles, 2017). RPE proteins were loaded into precast 4-12% NuPage Bis-Tris gels (Thermo Fisher/Life Tech, Carlsbad, CA) and resolved with PAGE. Proteins were transferred to PVDF membranes and the membrane blocked for 1 hr at RT in 5% dry milk, TBST (137 mM NaCl, 2.7 mM KCl, 19 mM Tris base with 0.1% Tween-20). The membranes were probed overnight at 4°C with antibodies dissolved in blocking buffer. We used the following antibodies for Western blotting: rabbit anti-catalase (Abcam ab1877, 1/5000), rabbit anti-Pex14 (ProteinTech 10594-1-AP, 1/1000), mouse anti-PMP70 (Sigma SAB4200181, 1/1000), rabbit anti-β-catenin (Cell Signaling, D10AB; 8480, 1/1000), mouse anti-β-actin (Sigma; A2228, 1/2500), and mouse anti-cyclophilin A (Cell Signaling; 2175S, 1/1000). Following washes, membranes were probed for 1 hr at RT with Horseradish peroxidase (HRP) - conjugated secondary antibodies were used to detect mouse (Thermo Fisher Scientific; G-21040, 1/2500) and rabbit IgG (ThermoFisher Scientific; 31460, 1/2500). Chemiluminescence signals were developed using SuperSignal® West Dura extended duration substrate (ThermoFisher Scientific; 34075).

### Catalase assays

Catalase activity was measured as amount of peroxide depleted in samples by detection with a probe reacting with H_2_O_2_ in the presence of HRP to produce the fluorescent resorufin (Amplex Red Catalase Assay Kit, Eugene, OR). Fluorescence at 590 nm was measured with a microplate reader (Fluoroskan Ascent microplate fluorometer, Thermo Systems). Catalase activity in experimental samples was determined as change in fluorescence compared to known standards. Lysates for measuring catalase activity were prepared similarly to Western blotting, except that solubilization was performed with manual homogenization in PBS. Triton X-100 was added to a final concentration of 0.5% followed by vigorous vortexing to enhance the extraction of catalase from peroxisomal membranes.

### Citrate synthase activity assays

Citrate synthase activity was measured using Mitocheck citrate synthase activity assay (Cat. # 701040, Cayman Chemical, Ann Arbor, MI.) as described (33). This assay is based on the change in absorbance measured with the release of SH-CoA due to the citrate synthase-catalyzed condensation of oxaloacetate and acetyl-CoA to form citrate. The reaction was initiated with the addition of oxaloacetate to reaction wells containing RPE lysates diluted ∼50-fold. Absorbance at 412 nm as a function of time was measured for lysate samples, positive controls (using a known amount of enzyme), as well as negative controls (with either no enzyme or no enzyme and no oxaloacetate). Specific activity was determined by dividing the reaction rate by lysate protein concentration.

## Acknowledgments

This work was supported in whole or in part, by the National Institutes of Health Grants EY010420-21 (to KBB), EY026525-02 (to KBB and NJP), R01EY027442 and P30EY000331 (to DSW), **and PENN Core grant.** The authors thank the PDM-live cell imaging core and the UPENN EM core for analyses.

## Abbreviations used

ABCD: ATP-binding cassette transporter subfamily D
ACOX: Acyl CoA Oxidase
AMD: age-related macular degeneration
βHB: β-hydroxybutyrate
DAB: diaminobenzidine
DHA: docosahexaenoic acid
FAO: fatty acid oxidation
HRP: Horseradish peroxidase
LC3B: microtubule-associated protein 1 light chain 3B
NBR1: neighbor of BRCA1 gene 1
OS: photoreceptor outer segment
P62: sequestosome 1; p62 protein
Pex14: peroxisomal biogenesis factor 14
PMP70: The ATP-binding cassette transporter subfamily D member 3
PPAR: peroxisome proliferator-activated receptor
RPE: retinal pigment epithelium
VLCFA: very long chain fatty acids

