## Supplemental Figure 1 for "Peroxisome turnover and diurnal modulation of antioxidant activity in retinal pigment epithelia utilizes microtubule-associated protein 1 light chain 3B"

### Supplemental Figure S1

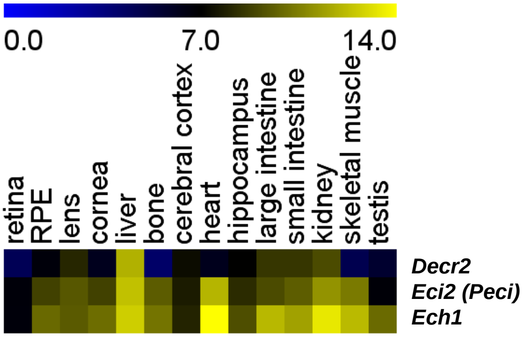


**Fig. S1 Relative expression of key peroxisomal enzymes required for completion of β-oxidation of DHA and other PUFA in various mouse tissues.** Microarray expression analysis using the database GSE10246 showing relative expression of *Decr2*: 2,4, Di-enoyl CoA reductase, *Eci2/Peci*: Peroxisomal Δ3,Δ2-enoyl-CoA isomerase, and Ech1: Δ3,5Δ2,4-dienoyl-CoA isomerase. Sequential activity of DECR2 and PECI is specifically required after the initial ACOX1- mediated desaturation of CoA esters of PUFA during the first round of β-oxidation in peroxisomes, and prior to subsequent β-oxidation reactions.
