## Supplemental Figure 2 for "Peroxisome turnover and diurnal modulation of antioxidant activity in retinal pigment epithelia utilizes microtubule-associated protein 1 light chain 3B"

### Supplemental Fig. S2

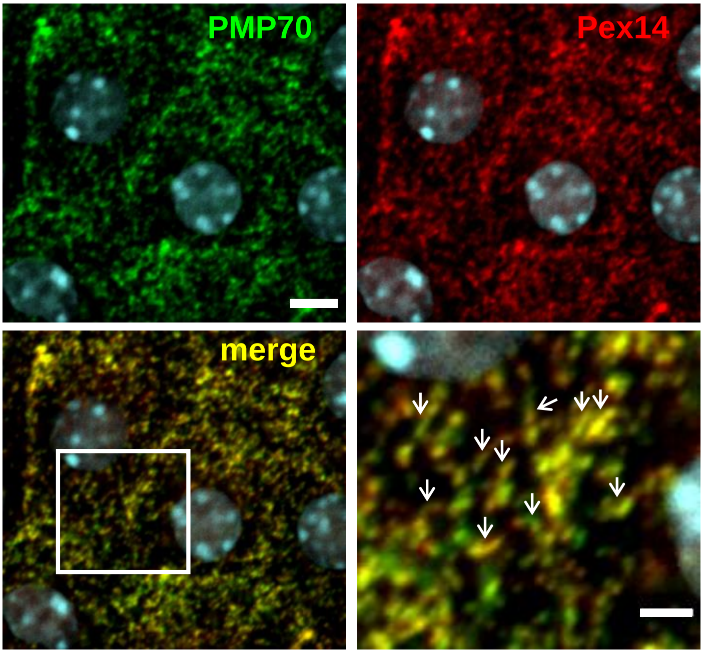


**Fig. S2 Peroxisome markers label tubular structures in mouse RPE** Confocal images at a single image plane of WT RPE/choroid/sclera bleached before immunolabeling with antibodies to PMP70 (green) and Pex14 (red). Lower right panel shows higher magnification view (boxed in lower left). Arrows denote examples of PMP70- and Pex14- co-labeled structures with a tubular appearance.
